# The effects of multifactorial stress combination on rice and maize

**DOI:** 10.1101/2022.12.28.522112

**Authors:** Ranjita Sinha, María Ángeles Peláez-Vico, Benjamin Shostak, Thi Thao Nguyen, Lidia S. Pascual, Sara I. Zandalinas, Trupti Joshi, Felix B. Fritschi, Ron Mittler

## Abstract

The complexity of environmental factors affecting plants is gradually increasing due to global warming, an increase in the number and intensity of climate change-driven weather events, such as droughts, heat waves, and floods, and the accumulation of different pollutants. The impact of multiple stress conditions on plants was recently termed ‘multifactorial stress combination’ (MFSC) and defined as the occurrence of three or more stressors that impact plants simultaneously or sequentially. We recently reported that with the increased number and complexity of different stressors, the growth and survival of *Arabidopsis thaliana* seedlings declines; even if the level of each individual stress is low enough to have no significant effect on plants. This finding is alarming since it reveals that MFSCs of different low-level stressors could impact crops and cause a dramatic reduction in overall growth. However, whether MFSC would impact commercial crop cultivars has not been studied. Here, we reveal that a MFSC of 5 different low level abiotic stresses (salinity, heat, the herbicide paraquat, phosphorus deficiency, and the heavy metal cadmium), applied in an increasing level of complexity, has a significant negative impact on the growth and biomass of a commercial rice (*Oryza sativa*) cultivar and a maize (*Zea mays*) hybrid. We further report on the first proteomics analysis of MFSC in plants that identified over 300 proteins common to all 4- and 5-MFSCs. Taken together our findings reveal that the impacts of MFSC on two different crop species are severe, and that MFSC may significantly affect agricultural productivity.

## INTRODUCTION

Global warming and climate change are subjecting plants to an increased frequency and intensity of different abiotic stressors that include droughts, heat waves, floods, and cold snaps (Bailey-Serres et al., 2019; IPCC 2021; Zandalinas et al., 2021a). In many instances these stressors occur together, for example during episodes of drought and heat waves (*e.g*., Mittler, 2006; Mittler and Blumwald, 2010; Zhang and Sonnewald, 2017; Alizadeh et al., 2020; Cohen et al., 2021). On top of these abiotic stresses and their combinations, are different man-made pollutants, such as heavy metals, microplastics, and pesticides, that affect plant growth and reproduction (Rillig et al., 2019, 2021; Zandalinas et al., 2021a). These could occur together with some of the different abiotic stresses and their combinations, highlighted above (Rillig et al., 2019, 2021; Zandalinas et al., 2021a; Zandalinas and Mittler, 2022). In addition to these stressors, are also climate-driven changes in the dynamics and distribution of different pathogen and insect populations that impact plants (Hamann et al., 2021; Kim et al., 2021). These conditions could be further augmented by poor nutrient content of different soils, as well as by a decrease in the complexity of soil microbiota. It was shown for example that the microbiome diversity of soils declined with the increased number of different climate change-driven stressors present in our environment (Rillig et al., 2019, 2021). The potential impact of the different complex abiotic and biotic stress conditions, described above, on plants was recently termed ‘multifactorial stress combination’ (MFSC) and defined as the occurrence of three or more different stressors that impact a plant simultaneously or sequentially (Rillig et al., 2019, 2021; Zandalinas et al., 2021a, 2021b).

We recently reported that with the increased number and complexity of different stressors, occurring together during a MFSC, the growth and survival of *Arabidopsis thaliana* seedlings declines; even if the level of each individual stress is low enough to have no significant effect on plants (Zandalinas et al., 2021a, 2021b). This finding is extremely alarming since it reveals that a MFSC of different low-level stressors (some already existing at different regions around the globe) could impact crops and cause a dramatic reduction in overall growth (Zandalinas and Mittler, 2022). However, whether MFSC would similarly impact commercial crop cultivars has not been reported. Here, we reveal that a MFSC of 5 different low level abiotic stresses (salinity, heat, paraquat, phosphorus deficiency, and cadmium), applied in an increasing level of complexity, has a significant negative impact on the growth and biomass of a commercial rice (*Oryza sativa*) cultivar and a maize (*Zea mays*) hybrid. To increase our understanding of the molecular mechanisms associated with MFSC, in a commercial cultivar, we conducted a proteomics analysis of rice seedlings subjected to MFSC. This analysis identified over 300 proteins common to all 4- and 5-MFSCs, including pathways involved in maintaining reactive oxygen species (ROS) and iron homeostasis. Taken together our findings reveal that the impacts of MFSC on two different crop species is severe and that MFSC may have a significant impact on agricultural productivity.

## RESULTS AND DISCUSSION

### Impact of MFSC on growth and biomass of rice and maize seedling

To determine whether MFSC would affect the growth and biomass of agricultural crops, we obtained seeds of a commercial rice cultivar (*Oryza sativa* var. Diamond), and a maize (*Zea mays* var. P1151AM) hybrid and studied their growth and biomass in response to MFSC. Seedlings were grown for 21 days under a combination of 5 different growth conditions (low-level stressors) that include: Salinity (50 mM NaCl), cadmium (400 μM CdCl_2_), paraquat (50 μM), heat stress (42°/36°C or 40°/32°C, day/night temperature, for rice or maize, respectively), and phosphorus deficiency. Control rice and maize plants were grown at 30°/26°C, day/night temperature. Each condition was applied individually and in all possible combinations, as previously reported for Arabidopsis (Zandalinas et al., 2021b). As shown in Figure 1, with the increase in the number and complexity of MFSCs, plant height, growth rate, and biomass of both rice and maize seedlings significantly declined. These findings suggest that while each of the different stresses had a non-significant effect on the rice and maize seedlings, when applied individually, the cumulative effect of all 4- or 5-low-level stressors significantly reduced rice and maize seedling’s growth and biomass (Figures 1, S1). Taken together, our findings reveal that, like Arabidopsis, commercial crop cultivars, in this case rice and maize, are negatively impacted by MFSC. This finding is important since Arabidopsis, which is extensively used in laboratory studies, is very different from commercial cultivars of crops used for agriculture (Mittler and Blumwald, 2010). In addition, while Arabidopsis is a dicot, rice and maize are monocots, showing that the negative effects of MFSC on plants are broad and can also apply to monocots. Although the different growth conditions used in this study may or may not occur in the natural or agricultural environment of different rice and maize genotypes, they nevertheless highlight the key principle of MFSC: With the increased number and complexity of different low-level stressors, occurring together during a MFSC, the growth of seedlings declines (Figure 1; Zandalinas et al., 2021a, 2021b; Zandalinas and Mittler, 2022).

**Figure 1.**
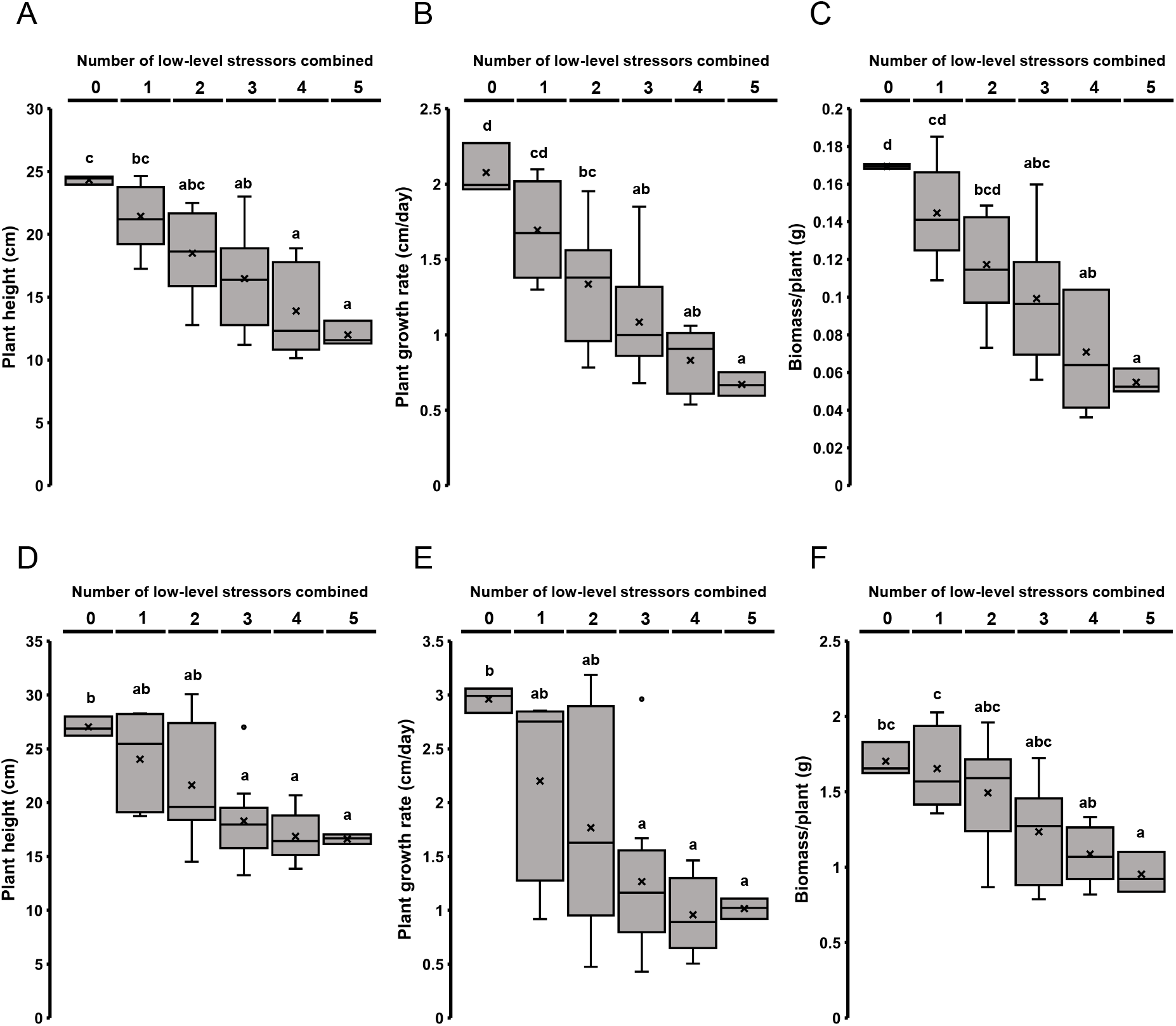
The impact of multifactorial stress combinations on the height, growth rate, and biomass of commercial rice (*Oryza sativa*) and maize (*Zea mays*) seedlings. The effects of multifactorial stress conditions (heat, salt, phosphorus deficiency, cadmium, and paraquat) applied in all different combinations (up to a combination of all five factors) was determined on the height, growth rate, and biomass of rice (**A-C**) and maize (**D-F**) seedlings. Box plots show the median (horizontal line), the lower and upper bounds of each box plot denote the first and third quartiles (the 25th and 75th percentiles, respectively), and whiskers above and below the box plot indicate 1.5 times the interquartile range. Statistical analysis was performed by one-way ANOVA followed by a Tukey post hoc test (different letters denote statistical significance at P < 0.05).

### Survival of rice seedlings under conditions of MFSC

To study the effects of MFSC on seedling survival in a crop plant, we focused on rice and used two different sets of growth conditions: (*i*) The MFSC conditions used above in Figure 1A (Figures 2A, S2A), and (*ii*) a different set of MFSC conditions that controlled for the possible interactions between paraquat and high light [Salinity (50 mM NaCl), cadmium (400 μM CdCl_2_), paraquat (50 μM), heat stress (40°/34°C day/night temperature), and low light (150 μmol photons m^-2^ s^-1^); Control plants were grown at 30°/26°C day/night temperature; 700 μmol photons m^-2^ s^-1^; Figures 2B; S2B]. The reason for using low light as a growth condition, was to control for the potential interactions between paraquat and high light intensities that can cause plant death (Zandalinas et al., 2021b). Low light intensity of 150 μmol photons m^-2^ s^-1^ is also considered a low-level stressor, as it provides rice plants with limited light energy for photosynthesis and growth (Yamori et al., 2016). As shown in Figure 2, each of the different individual growth conditions, as well as all the different 2- and 3-factor combinations, had no significant effect on the survival of rice. In contrast, the two different combinations of 5- low-level MFSCs had a significant effect on seedling survival, reducing it by about 40% (Figure 2), and the different 4-low-level stress combinations shown in Figure 2B had a significant effect of seedlings survival reducing it by about 25%. These findings suggest that while each of the different stress conditions used had a negligible effect on seedling survival, the cumulative effect of all 4- or 5-low-level stressors significantly reduced seedlings survival. Although our study was conducted with seedlings and did not evaluate grain yield per plant, a 25-50% decrease in biomass accumulation (Figure 1), and a 25-40% decrease in seedling survival (Figure 2), are likely to significantly decrease overall production of grain per unit area/field in rice.

**Figure 2.**
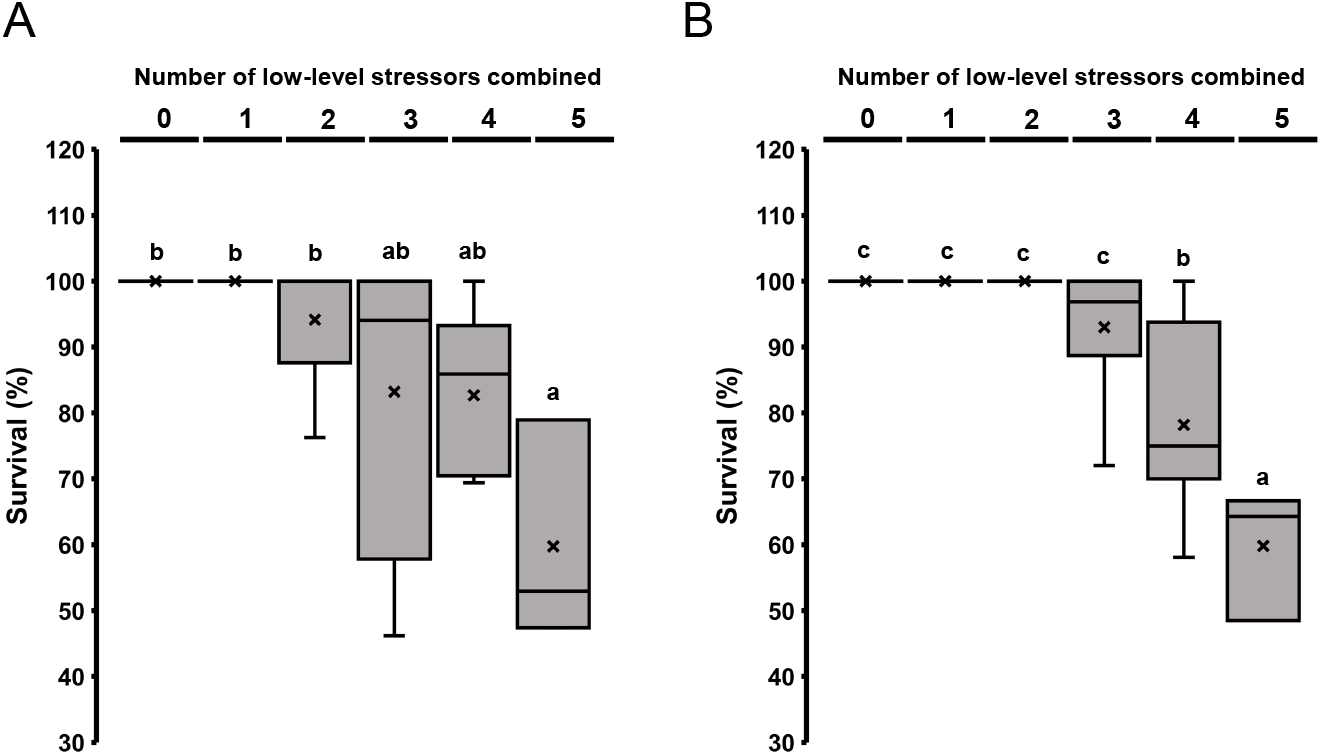
The impact of multifactorial stress combinations on the survival of rice (*Oryza sativa*) seedlings. The effects of multifactorial stress conditions (heat, salt, phosphorus deficiency, cadmium, and paraquat; **A**; or heat, salt, low light, cadmium, and paraquat; **B**) applied in all different combinations (up to a combination of all five factors) was determined on the survival of commercial rice seedlings. Box plots show the median (horizontal line), the lower and upper bounds of each box plot denote the first and third quartiles (the 25th and 75th percentiles, respectively), and whiskers above and below the box plot indicate 1.5 times the interquartile range. Statistical analysis was performed by one-way ANOVA followed by a Tukey post hoc test (different letters denote statistical significance at P < 0.05).

### Proteomics analysis of MFSC in rice seedlings

To gain better understanding of the molecular responses of a commercial cultivar to MFSC, and identify different proteins associated with MFSC in a crop plant (that could be used as an initial reference in breeding efforts), we conducted proteomics analysis of rice seedlings subjected to the MFSC shown in Figure 1A. As shown in Table 1, proteomics coverage was in the range of 4,100-4,600 identified proteins per treatment. To determine the effects of each stress, the abundance of the proteins identified in each treatment were compared to that of the control, and only proteins that had a significant change in their abundance (up- or down-regulated), compared to control in each treatment were considered for further analysis (Tables 1, S1-S31). Interestingly, under the conditions we used, paraquat and phosphorus deficiency resulted in a low number of proteins altered compared to control, (2 for paraquat and 8 for phosphorus deficiency). However, when these two stresses were combined (paraquat+phosphorus deficiency), 157 proteins were altered in their abundance. Similar findings were obtained with other individual low-level stresses and their combination [*e.g*., cadmium (78) and paraquat (2), and their combination (145), and cadmium (78) and phosphorus deficiency (8) and their combination (253); Table 1]. These findings suggest that with the increased complexity of some stresses, the response of plants increases, potentially indicating that some stresses may have a synergistic effect on each other (Zandalinas and Mittler, 2022). Of the different individual stresses used, heat stress had the highest impact on protein expression with over 1,500 proteins altered in their abundance (Table 1). This finding is consistent with the extensive impact of heat stress conditions on protein abundance in crops, including rice (*e.g*., Zou et al., 2011), and suggests that global warming is likely to play a key role in future responses of crops to MFSCs.

**Table 1.** Summary of proteomics results for the multifactorial stress combination analysis in rice.

| <u>Treatments</u> | <u>Proteins altered</u> | <u>Total number of proteins</u> |
| --- | --- | --- |
| CT |  | 4446.00 |
| S | 275 | 4600.00 |
| Cd | 78 | 4511.67 |
| Pq | 2 | 4472.33 |
| -P | 8 | 4483.67 |
| HS | 1520 | 4600.00 |
| S+HS | 1555 | 4541.67 |
| Cd+Pq | 145 | 4549.00 |
| -P+Cd | 253 | 4451.67 |
| Cd+HS | 1536 | 4424.33 |
| -P+Pq | 157 | 4315.00 |
| Pq+HS | 1515 | 4519.33 |
| S+Cd | 395 | 4468.33 |
| S+Pq | 575 | 4609.33 |
| -P+S | 622 | 4594.00 |
| -P+HS | 1472 | 4495.67 |
| S+Cd+HS | 1846 | 4483.67 |
| -P+S+Pq | 135 | 4330.67 |
| S+Pq+HS | 1609 | 4415.33 |
| -P+S+HS | 1628 | 4475.00 |
| -P+Cd+Pq | 232 | 4309.00 |
| -P+Cd+HS | 1651 | 4475.67 |
| S+Cd+Pq | 895 | 4478.67 |
| -P+S+Cd | 748 | 4500.67 |
| Cd+Pq+HS | 1630 | 4487.00 |
| -P+Pq+HS | 1536 | 4460.00 |
| -P+S+Cd+Pq | 1976 | 4103.67 |
| S+Cd+Pq+HS | 1208 | 4194.00 |
| -P+S+Pq+HS | 1505 | 4414.67 |
| -P+S+Cd+HS | 1739 | 4463.00 |
| -P+Cd+Pq+HS | 1318 | 4178.67 |
| -P+S+Cd+Pq+HS | 1402 | 4251.67 |

Average results are shown for each treatment (3 biological replicates). All proteins significantly altered in each treatment are shown in Supplementary Tables S1-S31. Abbreviations: CT, control; S, salt; Cd, cadmium; Pq, paraquat; -P, phosphorus deficiency; HS, heat stress.

To compare the overlap between the different stress treatments, we generated UpSet plots for all 2-, 3- (Figure S3), and 4- and 5-stress combinations (Figure 3A). This analysis revealed that the expression of 332 proteins was common to all 4- and 5-stress combinations, that resulted in the most severe impact on plant height, growth rate, biomass, and survival; Figures 1A, 2A). Gene ontology (GO) annotation term analysis of this group of proteins revealed that they were enriched in redox, catabolic metabolism, ROS scavenging, chaperone activity, iron-sulfur metabolism, and other functions (Figure 3B; see full list of GO terms in Table S32). To further study the abundance of ROS scavenging enzymes in our dataset, we generated a heatmap for the abundance of the key ROS scavenging enzymes ascorbate peroxidase (APX), catalase (CAT), glutathione reductase (GR) and superoxide dismutase (Mittler et al., 2022), found in all samples by our proteomics analysis (Figure 3C; Tables S1-S32). This analysis revealed that the abundance of many ROS scavenging enzymes (*e.g*., APX1, APX4, GR, and CAT-B) was elevated in samples from plants subjected to 4- or 5-stress combinations, compared to plants subjected to single stress conditions, or simple combinations of 2- and 3-stresses (Figure 3C). The identification of ROS and iron-sulfur metabolism categories in rice plants subjected to 4- and 5-stressors combined (Figures 3B, 3C) is in agreement with our previous transcriptomic study in Arabidopsis plants that identified these two categories as enriched in plants subjected to MFSC, as well as revealed that mutants deficient in ROS scavenging or signaling (*apx1* or respiratory burst oxidase homolog D; *rbohD*), or in balancing iron and ROS levels (plants with suppressed expression of AtNEET; AT5G51720) were less tolerant to MFSC (Zandalinas et al., 2021b). As ROS metabolism and NEET proteins, that regulate iron and ROS metabolism, are conserved among eukaryotic organisms (Mittler et al., 2019, 2022; Nechushtai et al., 2012), augmenting the ability of different crops to scavenge ROS or balance ROS and iron levels could be a viable strategy to increase their resistance to MFSC. Further studies are needed to determine the role of these pathways, as well as other pathways identified by our analysis (Figure 3; Tables S1–S32) in augmenting the tolerance of plants to MFSC. Our omics studies of MFSC in Arabidopsis (Zandalinas et al., 2021b) and rice (Figures 3, S3) further demonstrate that the response of plants to each different stress combination contains unique transcripts and proteins, and that only a few pathways are common to all different 4- and 5-stress combinations (*e.g*., ROS, iron metabolism, and chaperones). These could serve as a starting point in the search for genes that could augment the tolerance of different plants to different types of MFSCs (Zandalinas and Mittler, 2022; Rivero et al., 2022).

**Figure 3.**
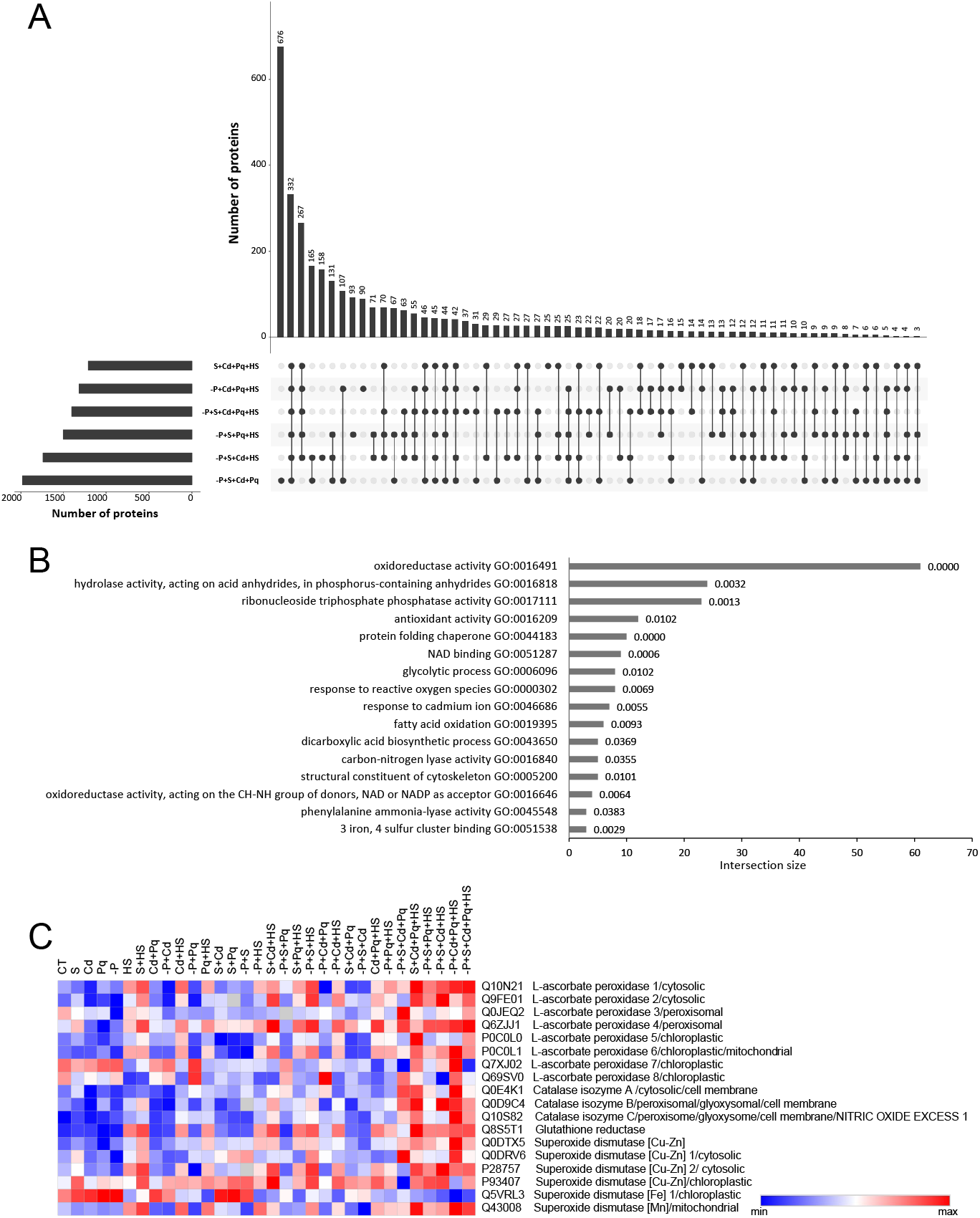
Proteomics analysis of multifactorial stress combination in rice seedlings. **A.** An UpSet plot showing the overlap between proteins significantly altered in their expression in all the different 4- and 5-stress combinations (UpSet plots for all 2- and 3-combinations are shown in Supplementary Figure S1). **B.** Selected gene ontology (GO) enrichment analysis terms for the 332 proteins common to all 4- and 5-combinations shown in A (a complete list of GO terms for the 322 proteins is shown in Supplementary Table S32). **C.** Heatmap for the abundance of different ROS-scavenging enzymes in all stress treatments used for the study. Benjamini-Hochberg with an FDR ≤0.05 was applied for proteomics analysis to determine significance. Summary of all proteomics results is shown in Table 1 and all proteins significantly altered in their abundance in each treatment are shown in Supplementary Tables S1-S31. Abbreviations: CT, control; S, salt; Cd, cadmium; Pq, paraquat; -P, phosphorus deficiency; HS, heat stress; ROS, reactive oxygen species.

### Conclusions

Our findings highlight the growing risk global warming, climate change, and industrial pollution pose to agriculture. The impact of MFSC on plants, documented under controlled environmental conditions (Figures 1, 2; Zandalinas et al., 2021b), highlight the urgent need for studies that quantify the impact of complex multifactorial stress factors under real-world conditions, as well as efforts aimed at identifying strategies to alleviate the impacts of MFSC on agriculture. Additional studies are also needed to determine the role of the different pathways identified by our proteomics (Table 1, Figure 3) and transcriptomics (Zandalinas et al., 2021b) analyses in augmenting the tolerance of different crops to MFSC. If we will not act to enhance the tolerance of our crops, and/or reduce the number, complexity, and intensity of different stressors affecting them, future episodes of MFSC could have a devastating impact on agriculture, potentially even leading to the destabilizing of multiple societies (Lobell et al., 2011; Challinor et al., 2014; IPCC 2021; Zsögön et al., 2021; Zandalinas et al., 2021a).

## MATERIAL AND METHODS

### Plant growth and stress treatments

All experiments were conducted in four identical growth chambers (BDR16, Conviron; Canada) under controlled growth conditions, and seedlings were randomized into the different growth conditions as described in (Sinha et al., 2022). Rice (*Oryza sativa* var. Diamond) and maize (*Zea mays* hybrid P1151AM) seeds, obtained from Tanner Seed Co., MO, USA, and Pioneer, Johnston, IA, USA, respectively, were germinated in peat and vermiculite growth media (1:1 mix) soaked in ¼ strength Hoagland solution with or without ammonium phosphate (Caisson Labs, Cat # HOP02, Smithfield, UT, USA), for control (CT) and phosphorus deficiency (-P), respectively. For CT conditions rice and maize seedlings were grown at 30/26°C - day/night temperature, 700 μmol photons m^-2^ s^-1^ light intensity, and 14/10 - hour day/night photoperiod. For salinity (S), cadmium (Cd), and/or paraquat (Pq) stresses, seeds were germinated in growth media mix soaked in ¼ Hoagland with 50 mM NaCl (Fisher Scientific, Hampton, NH, USA), 400 μM CdCl_2_ (Sigma-Aldrich, MO, USA), and/or 50 μM paraquat (Sigma-Aldrich, St. Louis, MO, USA). For salinity, cadmium, and/or paraquat, without phosphorus stresses, seeds were germinated in growth media mix soaked in ¼ Hoagland without phosphate with 50 mM NaCl, 400 μM CdCl_2_, and/or 50 μM paraquat. Briefly, 40 g of vermiculite and peat mix (1:1) was soaked with 160 ml of Hoagland solution (with/without stressors) and filled into free-draining pots of 12×8×6 cm^3^ dimension (length x width x height). About 20 or 12 rice or maize seeds were planted in each pot respectively (1 cm below soil surface). Pots were watered one more time with the above-mentioned stressors and their combinations. Afterwards, seedlings were periodically watered with deionized water and once a week with ¼ Hoagland with or without phosphate for the respective treatments, avoiding excessive watering. For CT, or stress treatments without HS, seedlings were grown at 30/26°C day/night temperature, 700 μmol photons m^-2^ s^-1^, 14/10 hour day/night photoperiod. For HS and the different combinations that included HS, rice plants were germinated and grown under 42/36°C day/night temperature, 700 μmol photons m^-2^ s^-1^, 14/10-hour photoperiod, while maize plants were germinated and grown at 40/32°C day/night temperature under the same light conditions. For low light (LL) stress conditions, rice seedlings were subjected to low light (150 μmol photons m^-2^ s^-1^, 30C/26°C day/night temperature, 14/10-hour photoperiod). Rice and maize MFSC experiments were carried out separately using the same growth chambers.

### Physiological measurements

Seedlings were grown for 21 days and scored for plant height at 10 and 18 days as described in Zandalinas et al., (2021b). Plant growth rate was calculated from the two height measurements of day 10 and 18. Survival, and shoot biomass were scored at day 21 as described in Zandalinas et al., (2021b) and Sinha et al., (2022), by weighing individual plants (shoot biomass) and scoring for the ability of plants to recover from the different stress treatment and re-grow (survival).

### Proteomics analysis

Rice shoots were collected at day 15 post germination from CT and all stress treatments (Table 1). About 20 rice shoots from each treatment were pooled and flash frozen in liquid nitrogen as an individual biological replicate, and the entire experiment contained 3 biological replicates per treatment. Total protein was isolated using the phenol extraction protocol described in (Mooney and Thelen, 2004), and protein pellets were resuspended in 6 M urea, 2 M thiourea and 100 mM ammonium bicarbonate. Protein concentration was determined using Pierce 660 nm Protein Assay (Thermo Fisher Scientific, Waltham, MA, USA). 30 μg of proteins from each sample were reduced, alkylated, and digested as described in Zandalinas et al., (2020). An EvoSep One liquid chromatography system coupled to a modified trapped ion mobility spectrometry quadrupole time-of-flight mass spectrometer (timsTOF Pro 2, Bruker Daltonik, GmbH, Germany) was used for all proteomics analyses as described by Zandalinas et al., (2020).

### Proteomics data analysis

The FragPipe computational platform (version 18.0) with MSFragger (version 3.5), Philosopher (version 4.4.0), and EasyPQP (version 0.1.33) components were used to build the spectral library, and the protein sequence database *Oryza sativa* subsp. Japonica, UniProt-UP000059680-48,899, was used for protein identification (da Veiga et al., 2020; Kong et al., 2017; Tyanova et al., 2016). dia-PASEF raw data was analyzed with DIA-NN version 1.8 (Demichev et al, 2020). Data was exported from DIA-NN for further analysis using Perseus version 1.6.15.0. Differential expression analysis by two-sided unpaired t-test was performed between each treatment and the CT (Zandalinas et al., 2020). Benjamini-Hochberg correction for multiple hypothesis testing was applied, with FDR ≤0.05 reported as significant. KEGG and quantification of significantly represented GO terms (q-value 0.05) were conducted using g:profiler (https://biit.cs.ut.ee/gprofiler/gost). Upset Plots were created in upsetr (https://gehlenborglab.shinyapps.io/upsetr).

### Statistical analysis

All experiments were conducted with 3 biological repeats. Each biological repeat was conducted with 3 technical repeats. Each technical repeat included 50 seedlings of rice or maize per treatment (20 rice seeds per repeat for proteomics). Treatments and growth chambers were randomized with each biological repeat. Statistical analysis for box plots in Figures 1 and 2 was performed using one-way ANOVA followed by Tukey post hoc test (different letters denote statistical significance at P < 0.05). Statistical analysis for Supplementary Figures S1 and S2 was performed using a student’s t-test (asterisks denote statistical significance at P < 0.05 compared to control).

## Acknowledgments

This work was supported by funding from the National Science Foundation (IOS-2110017, IOS-1353886, IOS-1932639), Interdisciplinary Plant Group, and University of Missouri.

## Author Contributions

R.S., M.A.P.V, T.T.N., L.S.P., and B.S. performed experiments and analyzed the data. R.M., F.B.F, R.S., T.J., M.A.P.V and S.I.Z. designed experiments, analyzed the data, and/or wrote the manuscript.

## Data Availability

The data that supports the findings of this study are available in the text, figure, and supplementary material of this article. Proteomics data was deposited in Pride (https://www.ebi.ac.uk/pride/), under the following accession number: PXD039065.

---

**Supplementary Figure S1.**
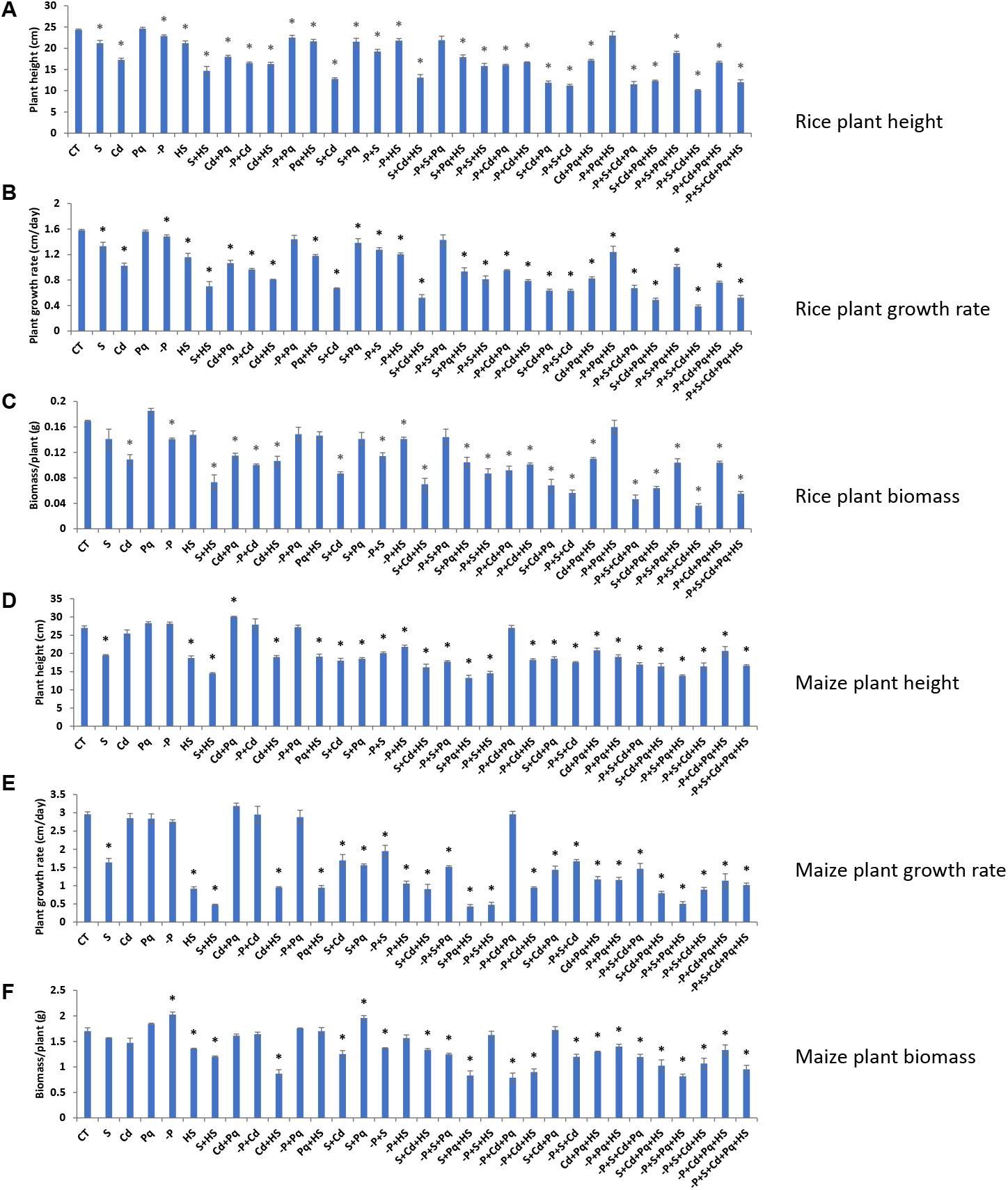
The impact of multifactorial stress combinations on the height, growth rate, and biomass of commercial rice (*Oryza sativa*) and maize (*Zea mays*) seedlings (In support of Figure 1). The effects of multifactorial stress conditions (heat, salt, phosphorus deficiency, cadmium, and paraquat) applied in all different combinations (up to a combination of all five factors) was determined on the height, growth rate, and biomass of rice (**A-C**) and maize (**D-E**) seedlings. Results are shown as average and SE for each treatment separately (significance change from control, *P < 0.05 was determined with a student’s t-test). Abbreviations: CT, control; S, salt; Cd, cadmium; Pq, paraquat; -P, phosphorus deficiency; HS, heat stress.

**Supplementary Figure S2.**
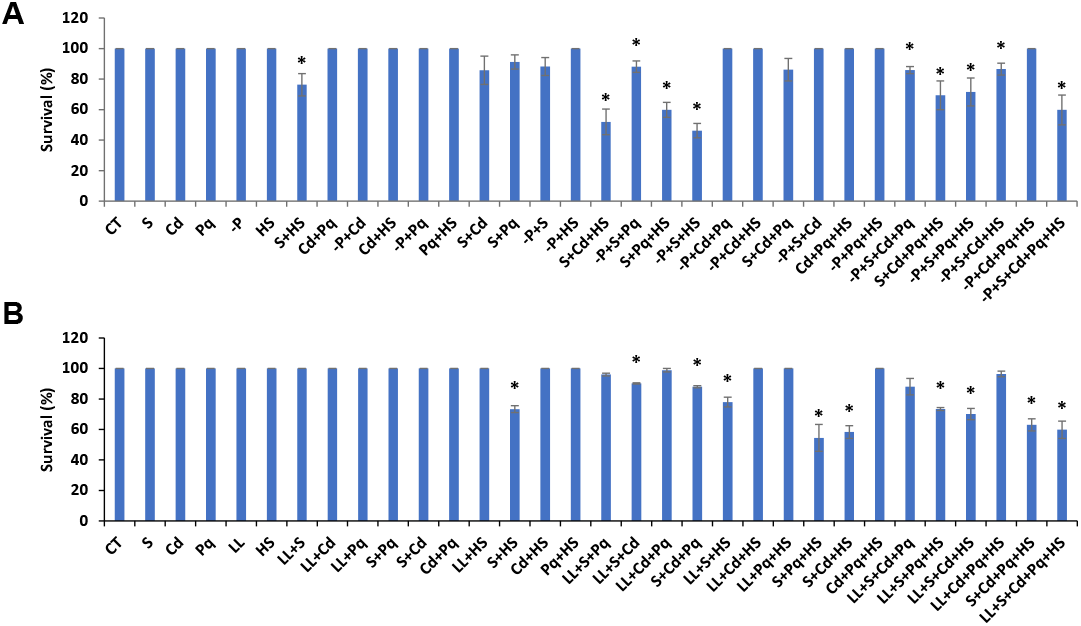
The impact of multifactorial stress combinations on the survival of rice (*Oryza sativa*) seedlings (In support of Figure 2). The effects of multifactorial stress conditions (heat, salt, phosphorus deficiency, cadmium, and paraquat; **A**; or heat, salt, low light, cadmium, and paraquat; **B**) applied in all different combinations (up to a combination of all five factors) was determined on the survival of commercial rice seedlings. Results are shown as average and SE for each treatment separately (significance change from control, *P < 0.05 was determined with a student’s t-test). Abbreviations: CT, control; S, salt; Cd, cadmium; LL, low light; Pq, paraquat; -P, phosphorus deficiency; HS, heat stress.

**Supplementary Figure S3.**
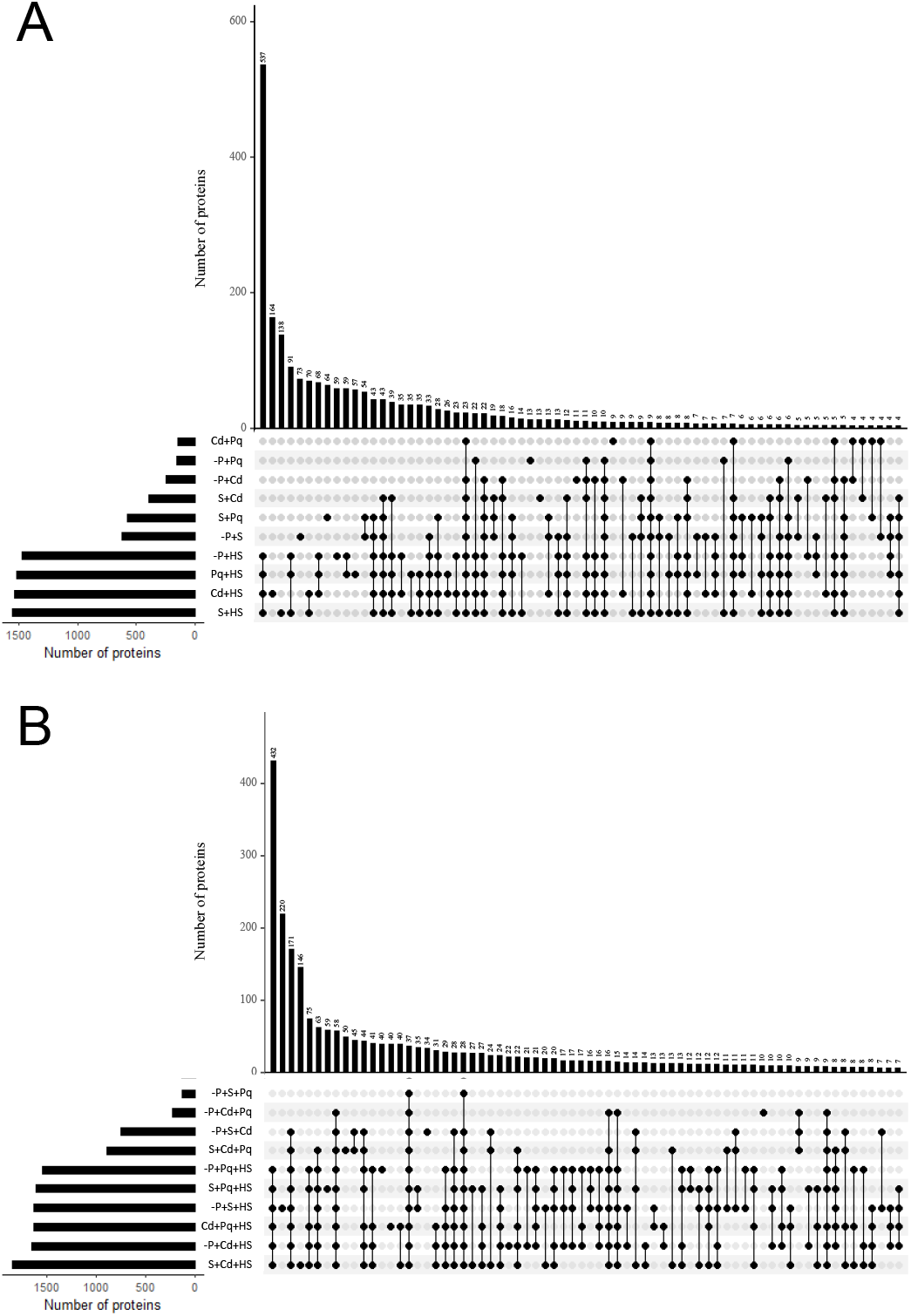
Proteomics analysis of multifactorial stress combination in rice seedlings (In support of Figure 3). **A.** An UpSet plot showing the overlap between proteins significantly altered in their abundance in all the different 2-stress combinations. **B.** An UpSet plot showing the overlap between proteins significantly altered in their abundance in all the different 3-stress combinations. Benjamini-Hochberg with an FDR ≤0.05 was applied for proteomics analysis to determine significance. Summary of all proteomics results is shown in Table 1 and all proteins significantly altered in each treatment are shown in Supplementary Tables S1-S31. Abbreviations: CT, control; S, salt; Cd, cadmium; Pq, paraquat; -P, phosphorus deficiency; HS, heat stress.

## Notes

### Competing Interest Statement

The authors have declared no competing interest.

## References

Alizadeh MR, Adamowski J, Nikoo MR, AghaKouchak A, Dennison P, Sadegh M (2020) A century of observations reveals increasing likelihood of continental-scale compound dry-hot extremes. Sci Adv 6: eaaz4571

Bailey-Serres J, Parker JE, Ainsworth EA, Oldroyd GED, Schroeder JI (2019) Genetic strategies for improving crop yields. Nature 575: 109–118

Challinor AJ, Watson J, Lobell DB, Howden SM, Smith DR, Chhetri N (2014) A meta-analysis of crop yield under climate change and adaptation. Nat Clim Change 4: 287–291

Cohen I, Zandalinas SI, Huck C, Fritschi FB, Mittler R (2021) Meta-analysis of drought and heat stress combination impact on crop yield and yield components. Physiol Plant. 171: 66–76

da Veiga Leprevost F, Haynes SE, Avtonomov DM, Chang HY, Shanmugam AK, Mellacheruvu D, Kong AT, Nesvizhskii AI (2020) Philosopher: a versatile toolkit for shotgun proteomics data analysis. Nat methods 17: 869–70

Demichev V, Messner CB, Vernardis SI, Lilley KS, Ralser M (2020) DIA-NN: neural networks and interference correction enable deep proteome coverage in high throughput. Nat methods 17: 41–44

Hamann E, Blevins C, Franks SJ, Jameel MI, Anderson JT (2021) Climate change alters plant–herbivore interactions. New Phytol 229: 1894–1910

IPCC 2021. Masson-Delmotte V, Zhai P, Pirani A, Connors S, Péan C, Berger S, Caud N, Chen Y, Goldfarb L, Gomis M, et al. (Eds.). Climate Change 2021: The Physical Science Basis. Contribution of Working Group I to the Sixth Assessment Report of the Intergovernmental Panel on Climate Change. UK: Cambridge University Press

Kim JH, Hilleary R, Seroka A, He SY (2021) Crops of the future: building a climate-resilient plant immune system. Curr Opin Plant Biol 60: 101997

Kong AT, Leprevost FV, Avtonomov DM, Mellacheruvu D, Nesvizhskii AI (2017) MSFragger: ultrafast and comprehensive peptide identification in mass spectrometry-based proteomics. Nat Methods 14: 513–520

Lobell DB, Schlenker W, Costa-Roberts J (2011) Climate trends and global crop production since 1980. Science 333: 616–620

Mittler R (2006) Abiotic stress, the field environment and stress combination. Trends Plant Sci 11: 15–19

Mittler R, Blumwald E (2010) Genetic engineering for modern agriculture: challenges and perspectives. Annu Rev Plant Biol 61: 443–462

Mittler R, Darash-Yahana M, Sohn YS, Bai F, Song L, Cabantchik IZ, Jennings PA, Onuchic JN, Nechushtai R (2019) NEET Proteins: A New Link Between Iron Metabolism, Reactive Oxygen Species, and Cancer. Antioxid Redox Signal 30:1083–1095

Mittler R, Zandalinas SI, Fichman Y, Van Breusegem F (2022) Reactive oxygen species signalling in plant stress responses. Nat Rev Mol Cell Biol 23: 663–679

Mooney BP, Thelen JJ (2004) High-throughput peptide mass fingerprinting of soybean seed proteins: automated workflow and utility of UniGene expressed sequence tag databases for protein identification. Phytochemistry 65:1733–1744

Nechushtai R, Conlan AR, Harir Y, Song L, Yogev O, Eisenberg-Domovich Y, Livnah O, Michaeli D, Rosen R, Ma V, et al. (2012) Characterization of Arabidopsis NEET reveals an ancient role for NEET proteins in iron metabolism. Plant Cell 24: 2139–54

Rillig MC, Lehmann A, Orr JA, Waldman WR (2021) Mechanisms underpinning nonadditivity of global change factor effects in the plant–soil system. New Phytol 232: 1535–1539

Rillig MC, Ryo M, Lehmann A, Aguilar-Trigueros CA, Buchert S, Wulf A, Iwasaki A, Roy J, Yang G (2019) The role of multiple global change factors in driving soil functions and microbial biodiversity. Science 366: 886–890

Rivero RM, Mittler R, Blumwald E, Zandalinas SI (2022) Developing climate-resilient crops: Improving plant tolerance to stress combination. Plant J 109: 373–389

Sinha R, Zandalinas SI, Fichman Y, Sen S, Zeng S, Gómez-Cadenas A, Joshi T, Fritschi FB, Mittler R (2022) Differential regulation of flower transpiration during abiotic stress in annual plants. New Phytol 235: 611–629

Tyanova S, Temu T, Sinitcyn P, Carlson A, Hein MY, Geiger T, Mann M, Cox J (2016) The Perseus computational platform for comprehensive analysis of (prote) omics data. Nat methods 13: 731–40

Yamori W, Makino A, Shikanai T (2016) A physiological role of cyclic electron transport around photosystem I in sustaining photosynthesis under fluctuating light in rice. Sci Rep 6: 20147

Zandalinas SI, Song L, Sengupta S, McInturf SA, Grant DG, Marjault HB, Castro-Guerrero NA, Burks D, Azad RK, Mendoza-Cozatl DG, Nechushtai R, Mittler R (2020) Expression of a dominant-negative AtNEET-H89C protein disrupts iron-sulfur metabolism and iron homeostasis in Arabidopsis. Plant J. 101: 1152–1169.

Zandalinas SI, Fritschi FB, Mittler R (2021a) Global warming, climate change, and environmental pollution: Recipe for a multifactorial stress combination disaster. Trends Plant Sci 26: 588–599

Zandalinas SI, Mittler R (2022) Plant responses to multifactorial stress combination. New Phytol 234: 1161–1167

Zandalinas SI, Sengupta S, Fritschi FB, Azad RK, Nechushtai R, Mittler R (2021b) The impact of multifactorial stress combination on plant growth and survival. New Phytol 230: 1034–1048

Zhang H, Sonnewald U (2017) Differences and commonalities of plant responses to single and combined stresses. Plant J 90: 839–855

Zou J, Liu C, Chen X (2011) Proteomics of rice in response to heat stress and advances in genetic engineering for heat tolerance in rice. Plant Cell Rep. 30: 2155–2165

Zsögön A, Peres LEP, Xiao Y, Yan J, Fernie AR (2021) Enhancing crop diversity for food security in the face of climate uncertainty. Plant J 109: 402–414

